# Hashimoto’s Thyroiditis-associated Thyroid Swelling among Adolescents Presenting with Benign Goiter in Lahore, Pakistan

**DOI:** 10.1101/468777

**Authors:** Saira Shan, Shumaila Aslam, Shan Elahi, Nayab Batool Rizvi, Nazish Saleem

**Author notes:** **Correspondence: Dr. Nayab Batool Rizvi**, Institute of Chemistry, New Campus, University of the Punjab, Lahore, Pakistan.

## Abstract

**Background:** The aim of this study was to determine frequency of Hashimoto’s thyroiditis (HT), by determination of serum TPO-Ab titer in goitrous adolescents attending Centre for Nuclear Medicine (CENUM), Mayo Hospital, Lahore, Pakistan.

**Method:** Serum TPO-Ab was determined in 277 local goitrous adolescents (female 194, male 83) with euthyroidism or subclinical hypothyroidism. Their mean (±SD) age was 15.8 + ***2.5 years with age range 10 to 19 year. Female and male adolescents were comparable in age, goiter size and serum thyroid hormone levels.

**Results:** Mean TSH and percentage of TSH levels > 3.0 mIU/L was significantly higher in male adolescents. High serum TPO-Ab titer (> 20.0 IU/ml) or HT was detected in 38 (13.7%) patients. The incidence of HT in female (16.5%) was higher than male adolescents (7.2%) but the difference was not statistically significant (p=0.120). Similarly goiter size (palpable or visible) or patient age (below or above 16 year) has no significant effect on HT frequency. However, compared to adolescents with TSH within normal laboratory range those with TSH level above the upper normal limit (4.0 mIU/L) had significantly more frequency of HT (10.4% versus 30.4%; p=0.001).

**Conclusion:** Thus this study reports 13.7% prevalence of HT-associated goiter among local adolescents presenting with benign goiter, in Lahore Pakistan.

## BACKGROUND

Thyroid gland in children and adolescents needs special care as thyroid hormones play important role in cellular metabolism, growth and development^1^. Puberty is a crucial period of complex hormonal interactions in which marked changes in thyroid function occur as an adaptation to body and sexual development^2^. That is why minimal diffuse enlargement of thyroid gland is found in many teenage boys and girls as a physiological response to such changes^1-5^. This enlargement usually regress^3^ but occasionally it may persist, enlarge and become nodular depending on many factors like gender, family history of thyroid disorder, iodine intake and presence of autoimmune thyroiditis (AIT)^2,6,7^. Low iodine intake enhances the TSH sensitivity and positive influence of growth factors involved in the physiological regulation of thyroid growth. The outcome of such stimulation may be substantial in girls leading to the development of goiter during mid to late puberty^2^. Such goiters are mostly colloidal and had been eliminated in many parts of the world after universal salt iodization (USI)^8, 9^. AIT is an inflammatory process and is the most common cause of thyroid enlargement and hypothyroidism in iodine sufficient areas^8, 10^. Thyroid enlargement due to AIT is termed Hashimoto’s thyroiditis (HT) ^10^.

In formerly iodine deficient areas where USI was successfully implemented emergence of HT in children and adolescents has been reported in the form of residual goiter ^8, 10, 11^. In countries like Pakistan where USI program is partially successful and is going towards success^12, 13^ etiology of an adolescent sporadic thyroid enlargement may either be iodine deficiency or AIT. A recent study conducted in Lahore reported that 21.2% of the adolescent girls had thyroid enlargement and majority of the goitrous girls (60%) were taking sufficient iodine^14^. Thus it seems that iodine deficiency is no more the sole cause of goiter and role of AIT may be suspected in the development of thyroid enlargement in local adolescents. The elucidation of exact etiology of an adolescent goiter is imperative because in contrast to iodine deficiency goiter thyroid enlargement due to AIT (HT) is a distinct entity. It is a complex disease that is also associated with other autoimmune diseases^5^. HT may lead progressively to worsening of thyroid function that had negative effects on growth and metabolic function in children and adolescents^4, 5^. This study was planned to know frequency of HT, by determination of serum TPO-Ab titer, in goitrous adolescents with euthyroid or subclinical hypothyroid function status detected at Centre for Nuclear Medicine (CENUM), Mayo Hospital Lahore, Pakistan.

## METHODS

CENUM, Mayo Hospital is one of the major referral centers for testing thyroid related disorders in the city and surrounding areas. We collected data of all referred goitrous adolescents, aged 10-19 years, who had undergone clinical assessment and determination of serum FT_4_, FT_3_ and TSH during calendar year 2015 and 2016. Adolescents with normal serum FT_4_ and FT_3_ (serum TSH normal or <10.0 mIU/L) were initially selected for this study. Adolescents already diagnosed with thyroid diseases and taking thyroid medications or had thyroid surgery were excluded. Similarly patients suffering from systematic diseases like DM, cardiac diseases and hepatitis were also excluded. Residual serum samples of finally selected adolescents, after testing thyroid related hormones were preserved for serum TPO-Ab determination. The Institutional Review Board at University of the Punjab, Lahore approved the study. A written consent for participation in study was obtained from each adolescent (age 18 year or above) or his/her parents (in case age under 18 year).

Initially a careful history of each patient was taken by the qualified physician. The physical examination of thyroid gland allowed assessment of goiter size by palpation. A 5 ml blood sample was drawn from each patient. The serum was separated by low-speed centrifugation (2000×g) for 5 minutes at room temperature. Serum samples were stored at −20°C until analysis. Serum samples were analyzed for FT_4_, FT_3_ and TSH. Serum FT_4_ and FT_3_ were estimated by radioimmunoassay (RIA) and TSH was estimated by imunoradiometric assay (IRMA) techniques using commercial kits of Immunotech Inc. (Beckman, Czech Republic). Serum TPO-Ab titer of residual serum sample was determined by ELISA method using commercial kit (IMMCO Diagnostics, Inc. NY, USA). Assay reliability was determined by the use of commercially derived control sera of low, medium and high concentrations which were included in every run. All assays were carried out in duplicate. Measurement of radioactivity, fitting of the standard curve and analysis of samples was carried out using a computerized gamma counter (Cap-RIA 16, CAPINTEC Inc. USA). RIA and IRMA results were expressed at less than 10% CV of imprecision profile. Normal ranges for serum FT_4_, FT_3_ and TSH, as standardized in our laboratory were 11.0 – 23.0 pmol/L, 2.8 – 5.8 pmol/L and 0.3 – 4.0 mIU/L respectively. The patients with TPO-Ab titer >20.0 IU/ml were considered positive according to instructions of kit manufacturer.

The analysis of data was carried out using Microsoft Excel program on a personal computer. Student T-Test and Chi-Square test was applied to test the significance of difference between two arbitrary groups. A value of p <0.05 was considered significant.

## RESULTS

A total of 277 adolescents presenting with goiter fulfilled the selection criteria and were selected for this study. All of them had serum FT_4_ and FT_3_ within respective normal ranges and TSH levels ranging 0.3 – 9.1 mIU/L. Among them 194 were female and 83 were male patients with average age 15.8 ± 2.5 year and age range 10 to 19 year. The anthropometric and laboratory characteristics of these adolescents as well as a comparison between male and female adolescents are shown in Table 1. Mean age and goiter sizes were comparable in male and female adolescents. Similarly mean levels of serum FT_4_ and FT_3_ were not significantly different between male and female participants. However, mean TSH level as well as percentage of serum samples with TSH > 3.0 mIU/L was significantly higher in male than female adolescents (p<0.05).

**Table 1.** - Anthropometric and laboratory characteristics of adolescents presenting with goiter

|  | All | Female | Male | P-value |
| --- | --- | --- | --- | --- |
| No. | 277 | 194 | 83 | - |
| Age (year) | 15.8±2.5 | 15.6± 2.8 | 15.9 ±2.3 | 0.140 |
| Grade 1 goiter, n (%) | 153 (55.2) | 100 (51.5) | 53 (63.9) | 0.165 |
| Grade 2 goiter, n (%) | 124 (44.8) | 94 (48.5) | 30 (36.1) | 0.165 |
| Mean FT<sub>4</sub> (pmol/L) | 16.5 ±2.9 | 16.6± 2.9 | 16.4 ± 2.7 | 0.533 |
| Mean FT<sub>3</sub> (pmol/L) | 4.5± 0.7 | 4.6 ± 0.8 | 4.7 ± 0.7 | 0.727 |
| Mean TSH (mIU/L) | 2.4± 1.9 | 2.2± 1.9 | 2.8 ± 1.9 | 0.018 |
| TSH ≥ 3.0 mIU/L, n (%) | 70 (25.3) | 39 (20.1) | 31 (37.4) | 0.008 |

Serum TPO-Ab titer above 10.0 IU/ml was detected in 46 adolescents ranging from 11.3 to 5904 IU/ml. Serum TPO-Ab titer higher than 20.0 IU/ml was detected in 38 (13.7%) adolescents. These adolescents were positive for TPO-Ab according to cutoff value provided by kit manufacturer. TPO-Ab positive adolescents as compared to TPO-Ab negative counterparts had comparable serum FT_4_ (16.0±3.2 vs 16.6±2.8; p=0.254) and FT_3_ (4.5±0.6 vs 4.7±0.8; p=0.457) but significantly higher TSH levels (3.7±3.1 vs 2.2±1.6; p=0.0062). The TPO-Ab positive adolescents covered the whole age range (10-19 year) and the youngest among them was 10 year old.

Table 2 shows the association of factors like gender, age, goiter size and serum TSH with positive TPO-Ab titer in adolescents. It was found that percentage of positive TPO-Ab titer was more than double in female than male adolescents but the difference was not significant (16.5% vs 7.2%; p=0.120). Similarly early to middle adolescent age group (age ≤ 16 year) had higher percentage of positive TPO-Ab cases than late adolescent group but difference was comparable (16.1% vs 10.7%; p=0.428). This difference remained same when only female adolescents were compared after same stratification (19.5% vs 12.3%; p=0.411). Also a non-significant difference was found between adolescents presenting with palpable and visible goiter. However, effect of serum TSH was decisive in this regard. The adolescents presenting with basal TSH level above upper limit of normal range (4.0 mIU/L) had significantly higher incidence of TPO-Ab positivity as compared to those with serum TSH within normal range (30.4% vs 10.4%; p=0.001). Further analysis showed that among subgroup of adolescents with normal TSH level female (n=155) and male (n=52) adolescents had non-significant difference of positive TPO-Ab titer (10.3% versus 5.8%; p=0.608) but this difference was significant between female and male adolescents with TSH level ≥ 3.0 mIU/L (41.0% versus 9.7%; p=0.014). Thus with overall HT incidence of 13.7%, female adolescents with TSH level above normal range had the highest percentage of HT among local goitrous adolescents.

**Table 2.** - Effect of gender, goiter size, age and basal serum TSH levelson HT incidence in goitrous adolescents

|  |  | TPO-Ab Positive,<br>n (%) | P-Value |
| --- | --- | --- | --- |
| Gender | Female (n=194) | 32 (16.5) | 0.120 |
|  | Male (n=83) | 06 (7.2) |  |
| Goiter Size | Palpable, G1 (n=153) | 21 (13.7) | 0.999 |
|  | Visible, G2 (n=124) | 17 (13.7) |  |
| Age | ≤16 Year(n=155) | 25 (16.1) | 0.428 |
|  | >16Year(n=122) | 13 (10.7) |  |
| TSH Level<br>(mIU/L) | Within NR (n=231) | 24 (10.4) | 0.001 |
|  | AboveNR (n=46) | 14 (30.4) |  |

NR = Normal Range

## DISCUSSION

Sporadic benign goiter is an enigma and debate on role of genetic or environmental factor in its etiology is still continuing^15^. In contrast to endemic goiter, etiology of benign sporadic goiter is complex. Besides iodine deficiency, thyroid autoimmunity, goitrogen ingestion, certain medicine and various infections may be involved in its etiology. It is very difficult to establish on clinical grounds alone which single or multiple factor is operative^16^. This is particularly true for detection of HT, the hallmark of which is the presence of high titer of serum TPO-Ab. The aim of this study was to know the prevalence of serum TPO-Ab in adolescents presenting with benign goiter so that contribution of autoimmune thyroiditis in etiology of thyroid enlargement may be documented. Our results showed that among male and female goitrous adolescents 13.7% have high serum TPO-Ab titer manifesting chronic autoimmune thyroiditis. Thus thyroid swelling (goiter) in these adolescents is not because of iodine deficiency but is a part of HT. This is probably the first large study of its sort conducted in goitrous adolescents residing in study area. A previous small study has detected high serum TPO-Ab titer in 19.5% goitrous adolescents mostly female^17^. Recently Ahmed et al reported HT in 9.7% of 73 thyroidectomy specimen collected from young goitrous patients (11-20 year) residing in hilly areas of Pakistan^18^. Thus ours and other local studies confirmed the emergence of HT as a cause of goiter in Pakistan. Our result is in accordance to Marwaha et al (1998) that reported HT in 13.8% of adolescent girls in India^8^. Comparable studies in other countries had reported 3-9% incidence of HT in children and adolescents presenting with goiter^11, 19, and 20^. The variation in prevalence of HT among different studies may be due to racial difference or different patient selection method employed in these studies.

The major risk factors for development of HT in adolescents are family history of thyroid disorder^21, 22^ and high iodine intake status^10, 20, 22^. Other contributory factors for HT are female gender^23, 24^, goiter size^22, 23^ and serum TSH level at presentation^23, 24^. We found that HT incidence in this study was independent of patient gender, age and goiter size and was positively associated only with serum TSH particularly above upper limit of laboratory range. Thus serum TSH was the only decisive factor in suspecting HT in local adolescents. These observations are in contrast to other studies reporting high incidence of HT in female patients^23, 24^ and among those presenting with large goiter^22^. The reason for TPO-Ab positivity in our patients may either be thyroid autoimmunity *per se* due to genetic component^25^, its induction after use of iodized salt for its treatment^20, 21^ or both. Enhanced dietary iodine content after successful implementation of USI had been implicated to initiate or aggravate the thyroid autoimmunity that cause thyroid swelling in children and adolescents^11, 20^. Increased levels of autoantibodies against thyroglobulin (Tg) and/or TPO had been detected in serum sample of goitrous children and adolescents taking iodized salt^10, 11, 20^. As information regarding iodine salt use in these adolescents is not available so no definite role of these factors may be ascertained for presence of high titer of TPO-Ab in these goitrous patients. However, a recently conducted study in female adolescents at our Centre revealed that 64.7% of HT patients were consuming iodized salt^26^. So role of excess iodine consumption in initiation or aggravation of autoimmune thyroiditis cannot be excluded.

Simple benign goiter is still a therapeutic challenge in medical practice. Use of iodized salt for goiter treatment may be beneficial to those having iodine deficiency induced goiter^27^ but for HT patients it is of no benefit. The continued use of iodized salt for HT treatment may possibly cause harm and lead to development of hypothyroidism. The treatment of choice for HT is thyroxine^28^. Along with this a careful supplementation of selenium and vitamin D in case of their deficiency is also recommended^29^. Thus differentiation of HT among adolescents presenting with benign goiter is necessary for proper treatment and follow up.

This study has many shortcomings. First, the number of male adolescents was low as compared to female. This has hampered the true comparison of relative incidence of HT in female and male adolescents. Second urinary iodine was not determined in adolescents to elucidate role of dietary iodine in HT. Moreover, data regarding family history of thyroid disorder was missing. A large study is warranted to investigate relative contribution of current iodine supplementation as well as family history of thyroid disorder in HT development in children and adolescents presenting with goiter.

## CONCLUSION

This study reports a prevalence of 13.7% of HT among local adolescents presenting with benign goiter. It may be speculated that emergence of HT is a natural consequence of USI started more than two decade ago in Pakistan. Further studies are required to characterize HT in different age groups of population. At present, realization of the presence and proper determination and treatment of this entity is necessary for thyroid heath in Pakistan.

## DECLARATIONS

### Abbreviations

AIT: autoimmune thyroiditis
TSH: thyroid stimulating hormone
USI: universal salt iodization
HT: Hashimoto’s thyroiditis
TPO-Ab: thyroid peroxidase antibodies
CENUM: Centre for Nuclear Medicine
FT_4_: free 3, 5, 3/, 5/-tetra-iodothyronine
FT_3_: free 3, 5, 37-triiodothyronine
ELISA: Enzyme linked immune sorbent assay
RIA: radioimmunoassay
IRMA: imunoradiometric assay
CV: cumulative variation
Tg: thyroglobulin
pmol/L: pico mole per litre
mIU/L: milli international unit

## Ethics approval and consent to participate

The Institutional Review Board at University of the Punjab, Lahore approved the study. A written consent for participation in study was obtained from each adolescent (age 18 year or above) or his/her parents (in case age under 18 year). [Mentioned in section “METHODS”]

## Consent for publication

All authors are agree to publish this study

## Availability of data and material

Data can be provided on demand by sending an email to

## Authors’ contributions

All authors participated in study. S.S (first author) and S.A (first co-author) conceived the idea of study, its plan of execution and collection of blood samples, S.E (second co-author) wrote the draft of the manuscript and analyzed the data, N.B. R (corresponding author) and NS (last author) carried out laboratory work and reviewed the manuscript.

## ACKNOWLEDGEMENTS

We are highly thankful to the doctors and Nurses and all the paramedic staff of Mayo Hospital, Lahore, Government Lady Atchison Hospital, Lahore, Pakistan and CENUM (Centre for Nuclear Medicine), Mayo Hospital Lahore, Pakistan for being supportive in collection of blood sample and data from the patients.

